# Patterns of gene family evolution and selection across *Daphnia*

**DOI:** 10.1101/2025.04.14.648758

**Authors:** Connor S. Murray, Alan O. Bergland

## Abstract

Gene family expansion underlies a host of biological innovations across the tree of life. Understanding why specific gene families expand or contract requires comparative genomic investigations clarifying further how species adapt in the wild. This study investigates the gene family change dynamics within several species of *Daphnia*, a group of freshwater microcrustaceans that are useful model systems for evolutionary genetics. We employ comparative genomics approaches to understand the forces driving gene evolution and draw upon candidate gene families that change gene numbers across *Daphnia*. Our results suggest that genes related to stress responses and glycoproteins generally expand across taxa, and we investigate evolutionary hypotheses of adaptation that may underpin expansions. Through these analyses, we shed light on the interplay between gene expansions and selection within other ecologically relevant stress response gene families. While we show generalities in gene family turnover in genes related to stress response (i.e., DNA repair mechanisms), most gene family evolution is driven in a species-specific manner. Additionally, while we show general trends towards positive selection within some expanding gene families, many genes are not undergoing selection, highlighting the complex nature of diversification and evolution within *Daphnia*. Our research enhances the understanding of individual gene family evolution within *Daphnia* and provides a case study of ecologically relevant genes prone to change.

## Introduction

A major goal in biology is to understand how adaptive evolution changes complex phenotypes (Lewontin, 1974; Mayr, 1963). Gene family expansion, mediated through the novel duplication of genes, is an influential process that provides species with the opportunity for biological innovation to occur (Hahn et al., 2007; Jordan et al., 2001). Neofunctionalization is the process wherein gene copies adopt new function (Ohno, 2013), while subfunctionalization is the process in which a paralogous gene retains a part of the progenitor’s role (Lynch & Force, 2000). Both neofunctionalization and subfunctionalization are ways in which gene duplications lead to diversification resulting from the fixation of beneficial amino acid substitutions (Lynch, 2002). Gene family expansions have long been hypothesized as the product of adaptive evolution across the tree of life, from microbes to mammals (Huang et al., 2023; Zhang et al., 2014; Hahn et al., 2007; Jordan et al., 2001; Lugli et al., 2017; Richter et al., 2018).

When developing hypotheses for studying gene family evolution we often view gene content change as facilitating adaptation to environmental (abiotic and biotic) shifts. For instance, animals inhabiting extreme temperature regimes typically have an increased number of heat-shock proteins (Chen et al., 2018; Zhang et al., 2012). In this way, gene family expansions have allowed taxa to adapt and survive during temperature fluctuations (Lindquist & Craig, 1988). While heat-shock proteins are known candidates of environmentally-induced gene family expansions, opsins are another source of constant gene family fluctuations, due to adaptation to various light environments and ultra violet sensitivities (Novales Flamarique, 2013), as well as chemosensory gene families (Peñalva-Arana et al., 2009). Additionally, innate immune response proteins and DNA repair mechanisms are also common expansion targets (Teekas et al., 2022), yet lineage specific expansions are also expected due to specific adaptation to local environments (Lespinet et al., 2002). Overall, interpreting how gene family changes occur across related species is a worthwhile pursuit, especially for taxa prone to gene family turnover in response to environmental decay because it reveals a component of adaptation that changes many potential protein targets across an organism (Guijarro-Clarke et al., 2020). By using comparative genomics, we can test hypotheses related to the timing of gene family diversification and lay the groundwork for better understanding adaptation within the context of both individual species and groups of taxa (Mendes et al., 2021).

In this work, we assess gene family evolution and infer the strength of selection acting upon *Daphnia*, a diverse genera of small-freshwater Crustaceans that live in a range of habitats from ephemeral rain-puddles to lakes and even manmade estuaries (Fryer 1991). *Daphnia* adaptively radiated roughly 200 million years ago (mya) and encompasses at least 121 species to date within the Daphniidea family. Subspecies and cryptic speciation is common within *Daphnia* and so this number of species is likely an underestimate (Forró et al., 2008). One of the most studied taxa within Daphniidea is *Daphnia pulex*, a complex of cryptic species found across both North American and European ponds (Colbourne et al., 1998; Crease et al., 2012; Murray et al., 2024; Vergilino et al., 2011). The first Crustacean genome described was *D. pulex* (Colbourne et al., 2011) and subsequent studies showed that even within species’ lineages of *D. pulex* shows variability of the number of genes present (Brandon et al., 2017; Lynch et al., 2017).

*Daphnia* species show fluctuations in the spectrum of gene gain and loss in response to environmental cues, supporting the case that gene family change is an important evolutionary mechanism in these taxa (Hamza et al., 2023; Schurko et al., 2009; Zhang et al., 2021). The focal ponds that *Daphnia* inhabit will periodically go through environmental degradation and hypoxia (Paul et al., 1998). Occasionally this hypoxia is driven by eutrophication caused by runoff of nutrients into freshwater systems (Ebert, 2022; Frisch et al., 2014). While this environmental stress largely occurs in areas devoted to agriculture, it can nonetheless affect species across entire ranges, especially in areas prone to temperature fluctuations (Smith & Schindler, 2009). One way in which *Daphnia* responds to eutrophication-induced hypoxia is to increase production of hemoglobin, which will aid oxygen binding (Fox et al., 1951). This increase in heme production results in a body-wide red hue that is potentially conserved across *Daphnia* (Zeis, 2020).

*Daphnia* are also notable because they have a unique reproductive mode of cyclical parthenogenesis whereby females have rounds of clonal reproduction followed by a sexual event triggered through environmental cues (Rouger et al., 2016). Cyclical parthenogenesis is common across the tree of life, yet we only have a limited understanding of the gene family pathways leading to this reproductive novelty, of which meiotic pathways are thought to be especially important (Schurko et al., 2009). Because some *Daphnia* express both asexual and sexual modes, their molecular machineries to accommodate these phenotypes must be conserved in the same genome, making them an attractive model to understand reproductive-linked gene evolution (Lumer, 1937; Zaffagnini & Sabelli, 1972). Outside of reproductive mode variation, sperm morphology is highly polymorphic and is divergent in tail length between the *Daphnia* and *Ctenodaphnia* subgenera (Duneau et al., 2022). Sperm morphology differences motivates the question of which genes have expanded or contracted across *Daphnia* and *Ctenodaphnia*? And are there ecologically relevant gene families related to stress, immune responses, and reproduction that have evolved across *Daphnia*?

In this work, we survey gene family evolution across *Daphnia*, highlight the expanding gene families, and we measure the strength of natural selection acting upon candidate genes. In this way, we test an overarching hypothesis that expanding gene families are also under positive selection. Our results show substantial gene content shifts across species and that stress response (DNA repair) and glycoprotein associated gene families largely fluctuate across *Daphnia* genomes. We detect positive selection within some of these overrepresented gene families, indicating a link between selection and gene content expansion among ecologically relevant genes.

## Materials and Methods

### Daphnia whole-genome dataset

Chromosome and scaffold-level assemblies of seven species from the genus *Daphnia* were collected from the NCBI Genome search engine (https://www.ncbi.nlm.nih.gov/datasets/genome/) accessed in January 2025 (Kitts et al., 2016). We chose North American *D. pulex* (KAP4; RefSeq: GCF_021134715.1), European *D. pulex* (D84A; GenBank: GCA_023526725; Barnard-Kubow et al., 2022), North American *D. pulicaria* (RefSeq: GCF_021234035.1; Wersebe et al., 2023), *D. sinensis* (GenBank: GCA_013167095.2, Jia et al., 2022), *D. carinata* (RefSeq: GCF_022539665.2), *D. galeata* (GenBank: GCA_030770115.1; Nickel et al., 2021), and *D. magna* (RefSeq: GCF_020631705.1) for analyses because they are the most complete species representatives and were annotated for protein-coding genes. and these genomes were the highest quality and newest available for each unique species. We also included *Artemia franciscana* (i.e., brine shrimp; GenBank: GCF_032884065.1) and *Penaeus monodon* (i.e., black tiger shrimp; GenBank: GCF_015228065.2) as Crustacean outgroups for our tree-based analyses.

### Computational biology pipeline

We implemented a *Nextflow v23.04.1* (Di Tommaso et al., 2017) pipeline (named *nf-GeneFamilyEvolution*) to organize the “flow” of data between processes and implement reproducibility and accessibility across computational environments. Briefly, we set up the nextflow pipeline to take RefSeq/GenBank genome identifiers, download their genome and proteome from the NCBI, classify ortholog groups, run various software related to phylogenetic tree building, and preform gene family evolution analyses with minimal setup and editing (Supplemental Figure 1A). The following sections describe individual components that are incorporated in the pipeline.

**Supplemental Figure 1:**
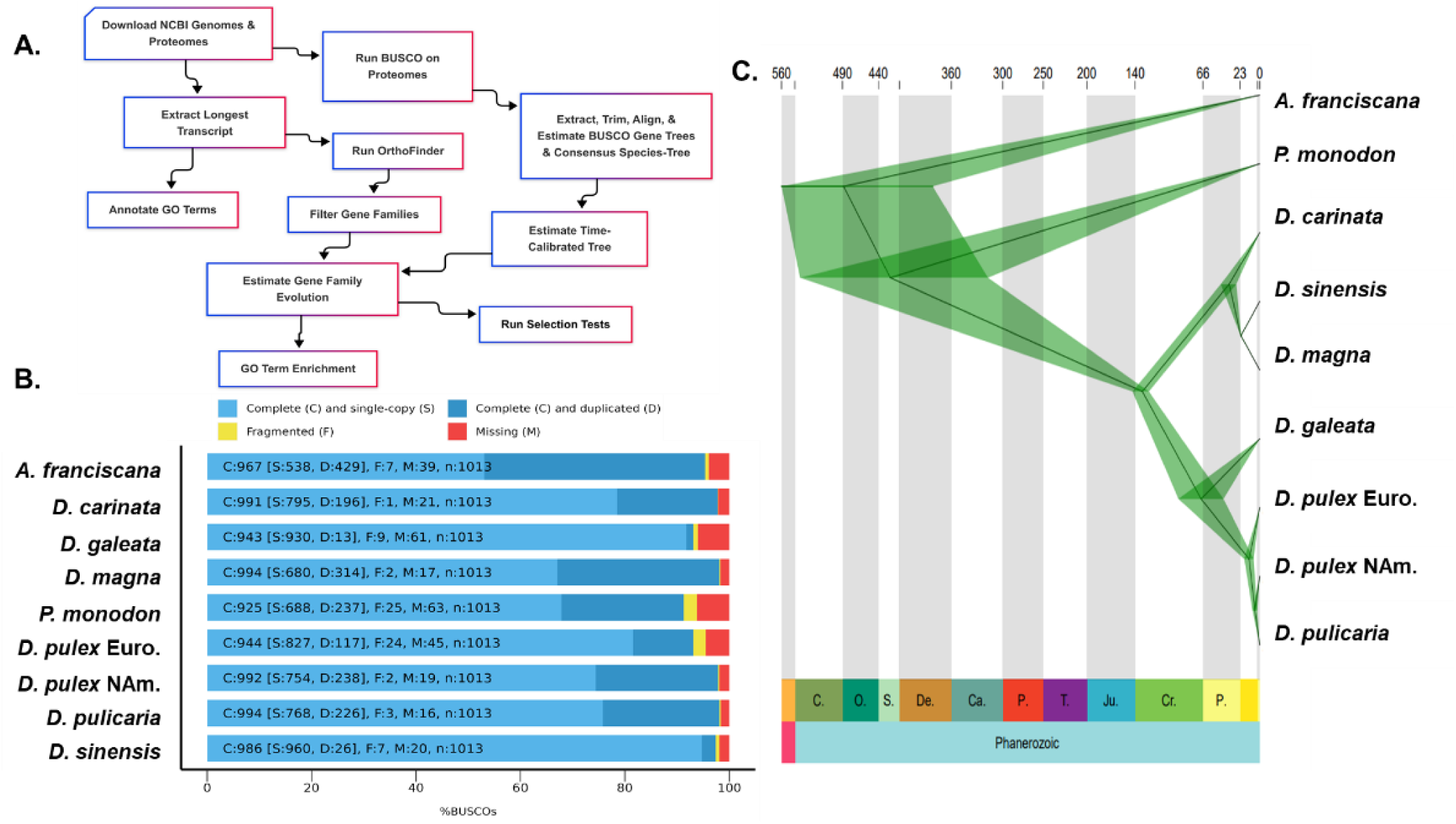
Pipeline and BUSCO Species tree. **A)** Nextflow pipeline showing the flow of data between processes starting with a list of RefSeq/GenBank identifiers**. B)** BUSCO scores denoting the number of genes that are complete, single copy, duplicated, fragmented, missing, and the total across each of the genomes against the Arthropoda dataset. **C)** Time-calibrated phylogenetic tree of the whole-genomes. The tree was built with the BUSCO genes that were complete and present within each genome. We ran each tree for five million Markov chain Monte Carlo generations.

### Estimating divergence-time across species

We used *BUSCO v5* (Manni et al., 2021) Arthropoda dataset on each genome assembly to acquire the benchmark universal single copy orthologous (BUSCO) score and used 164 complete single-copy genes for phylogenomic analyses of divergence dating across the *Daphnia* genera (Supplemental Figure 1B). To do this, we extracted each amino acid alignment for each gene using *seqkit faidx v2.2.0* (Shen et al., 2016) and aligned them using *mafft –auto v7.505* (Katoh & Standley, 2013). And then used *clipkit -m smart-gap v2.1.1* (Steenwyk et al., 2020) to clip out regions with large gaps. After this, we concatenated all of the protein sequences together using *seqkit concat v2.2.0* and used the *mcmctree_tree_prep.py* script (https://github.com/kfuku52/mcmctree_tree_prep) to create the necessary input files for *MCMCtree v4.9e* (Dos Reis & Yang, 2019), which is part of the *PAML* package of software (Yang, 2007). We assembled a gene-tree using *IQtree v2.2.0.3* with the *modelfinder* option (Kalyaanamoorthy et al., 2017).

We used several time-calibration points from previous phylogenetic investigations in *Daphnia.* Specifically, we used *D. carinata* - *D. magna* [100.4 - 104.8 mya] in place of the subgenus *Daphnia - Ctenodaphnia* root comparison in Cornetti et al., (2019). *Artemia* and *D. magna* [365.1 – 492 mya], *Artemia* and *Penaeus* [275 – 541 mya], and North American *D. pulex* – *D. magna* [130 - 150 mya] were taken from *Timetree5* (Kumar et al., 2022; Mathers et al., 2013), and *D. magna* - *D. sinensis* [21.5 - 22.4 mya] was from Cornetti et al., (2019). We used the *MCMCtreeR v1.1* package in *R* to plot the 95% confidence interval of divergence estimates across taxa (Supplemental Figure 1C; Puttick, 2019). *MCMCtree* was run several times to ensure model convergence by showing minor deviations in the estimate of node divergence times and the mean time for each node.

### Gene family evolution analyses and ontology enrichment

We first identified and retained the longest transcript with an open-reading frame in the sequence for each gene using the *primary_transcript.py* script within the *OrthoFinder v2.5.5* tool set (Emms & Kelly, 2019). We then classified orthologous genes between the FASTA format proteomes using *OrthoFinder*. After identifying the orthologous genes between the seven species, we annotated the phylogenetic hierarchical orthologous groups (HOGs) with the most common gene name by a majority vote using the *annotate_orthogroups* function in *orthofinder-tools* (https://github.com/MrTomRod/orthofinder-tools). We used the HMMER dataset and *hmmer2go v0.18.2* functions (https://github.com/sestaton/HMMER2GO) to assign gene ontology (GO) terms to the HOGs for use in enrichment analyses. We preformed quality control by hand for some single-copy ortholog genes families identified in *OrthoFinder* by BLASTing each amino acid sequence against the NCBI database using *blastp v2.13.0* (Sayers et al., 2022). This quality control step verified the GO term assignments and supported the translated function of each protein-coding gene tested (N=15).

*CAFE5 v1.1* (Mendes et al., 2021) was used to estimate the expansions and contractions of gene families across the *Daphnia* proteomes. Before running *CAFE5*, we excluded any HOGs that had over 80 genes present within any one species and any genes that were exclusively present in only one species, according to best practices in the *CAFE5* vignette. After this, we ran different models and found that the model with a varying gamma rate and root Poisson distribution (-k3 -p) fits our dataset and converged. After this, we extracted the HOGs found to be significantly expanding or contracting within each species and used those genes as the foreground and each species’ genome as the background with *clusterProfiler v3.14.3* in *R* (Wu et al., 2021). We used *REVIGO v1.8.1* as a semantic reduction tool to minimize GO term redundancy for any enriched terms (Supek et al., 2011), and preformed Bonferroni-Holm multiple testing corrections on *p*-values using *p.adjust* in *R*.

### Hypothesis testing of positive selection on codon sequences

To test for positive selection across gene families, we used *hyphy v2.5 aBSREL* (Kosakovsky Pond et al., 2020) on aligned codon FASTAs generated from *OrthoFinder*. *aBSREL* tests for positive selection occurring across branches of a tree by varying the rate of selection *dN/dS* (*ω*) across both sites and branches, in this way it models both site- and branch-level *dN/dS* heterogeneity (Smith et al., 2015). *aBSREL* fits a full adaptive model, a likelihood ratio test is then preformed at each branch and compares the full model to a null model where branches are not allowed to have rate classes of *dN/dS* > 1 (Kosakovsky Pond et al., 2020). As recommended by tutorials on *aBSREL*, we preformed one test on each tree, comparing all leaf nodes (i.e., tips) in a pairwise manner (Spielman et al., 2019). And tested trees to understand the patterns of selection occurring on those that have potentially undergone neofunctionalization (Hou et al., 2013; Mulhair et al., 2023; Saad et al., 2018; Wang et al., 2023). We translated the amino acid alignments and corresponding nucleotide sequences with *pal2nal.pl v14* (Suyama et al., 2006) and excluded any premature stop codons and gaps. We next aligned the sequences with *mafft* and visually inspected alignments for quality and removed any alignments that had evidence of artificial frameshifts. We ran each translated codon FASTA independently with ten cores and fixed the gene-tree for usage in each *aBSREL* run. We read the *aBSREL* json files into *R* for further analysis and plotting. We also used the *Datamonkey v2.0* webserver (Weaver et al., 2018) to export trees from the *aBSREL* models.. We counted a gene family as being under significant positive selection where the 1 < *dN/dS* (*ω*) < 10 and the multiple-testing corrected *p*-value < 0.05. We tested the odds ratio enrichment of the gene families that belong to the expanded genes identified from *CAFE5* against the genes that are non-fluctuating using a two-tailed Fisher’s exact test in *R*.

### Statistics and visualization in R

Statistical analyses were preformed using *R v4.3.1* (R Core Development Team). We used the following *R* packages for general analysis and visualization: *tidyverse v1.3.1* (Wickham et al., 2019), *ggplot2 v3.3.5* (Villanueva & Chen, 2019), *ggtree v2.0.4* (Xu et al., 2022), *patchwork v1.0.1* (Pedersen, 2022), *data.table v1.12.8* (Dowle & Srinivasan, 2023), *foreach v1.4.7*, *doMC v1.3.5* (Daniel et al., 2022).

### Data accessibility statement

All scripts and data used in every analysis are deposited on our GitHub repository along with the nextflow pipeline: https://github.com/connor122721/nf-GeneFamilyEvolution. All relevant data are deposited on data dryad at DOI:10.5061/dryad.gqnk98t02.

## Results

### Daphnia genomes and the gene family dataset

From the *Daphnia* genomes, we first identified 160,498 unique genes across the whole dataset after extracting the longest open reading frame per protein coding transcript. From these, *Orthofinder* identified 1,129 single-copy orthologous gene families. Within the total gene dataset, *Orthofinder* found 13,784 hierarchical phylogenetic orthologous gene groups (HOGs). We use these 13,784 HOGs as input into *CAFE5* to estimate the evolutionary rate of gene family gain and loss and to identify any gene groups that are evolving across the phylogeny. Below, we use this gene grouping information to expand our understanding of the phylogenetic relationship between taxa.

### Phylogeny of the represented Daphnia genomes

We built a time-calibrated phylogenetic tree to understand the relatedness of each *Daphnia* genome as well as create a phylogeny to serve in our gene family analyses (Supplemental Figure 1C). We find that North American and European *D. pulex* diverged 12.5 million years ago (mya) [95% confidence intervals; 7.2, 18.2]. Recent work highlights a split at 10 mya using BUSCO gene SNPs, well in range of our estimates (Murray et al., 2024). Previous estimates of the split time for North American and European *D. pulex* place it as early as 2-3 mya based on mitochondrial genes and assorted DNA loci (Colbourne et al., 1998; Crease et al., 2012). We also show that North American *D. pulex* and *D. pulicaria* diverged 5.6 mya [2.9, 8.5], an estimate higher than pervious at 0.5 - 2 mya (Crease et al., 2012). We also refine the split between the *Ctenodaphnia* and *Daphnia* subgenera to 35.8 mya [27.3, 45.5] (Supplemental Figure 1C).

### Trends of gene family evolution

We used *CAFE5* to identify expanding and contracting gene families across our dataset (Figure 1; Mendes et al., 2021). The *k3p CAFE5* model minimized the negative log-likelihood value, so we are reporting its output. Our first finding is that *D. magna* has the largest expansion within their genome (N_Genes_ = 1,054; Figure 1), while European *D. pulex* has the largest contraction (N_Genes_ = 1,309; Figure 1). Also, across the dataset, there are more expansions than contractions when taking the average loss and gain across individual genomes. There are 673 individual genes being lost and 567 genes being gained. And the range in gene gains [202-1,054] and loss [249-1,309] are wide for both classes. For the remainder of our work, we investigate the genes belonging to the 3,161 phylogenetic hierarchical ortholog groups (HOGs) identified as being significant candidates that are evolving across the species tree as well as ecologically relevant gene families related to stress response are tested below.

**Figure 1:**
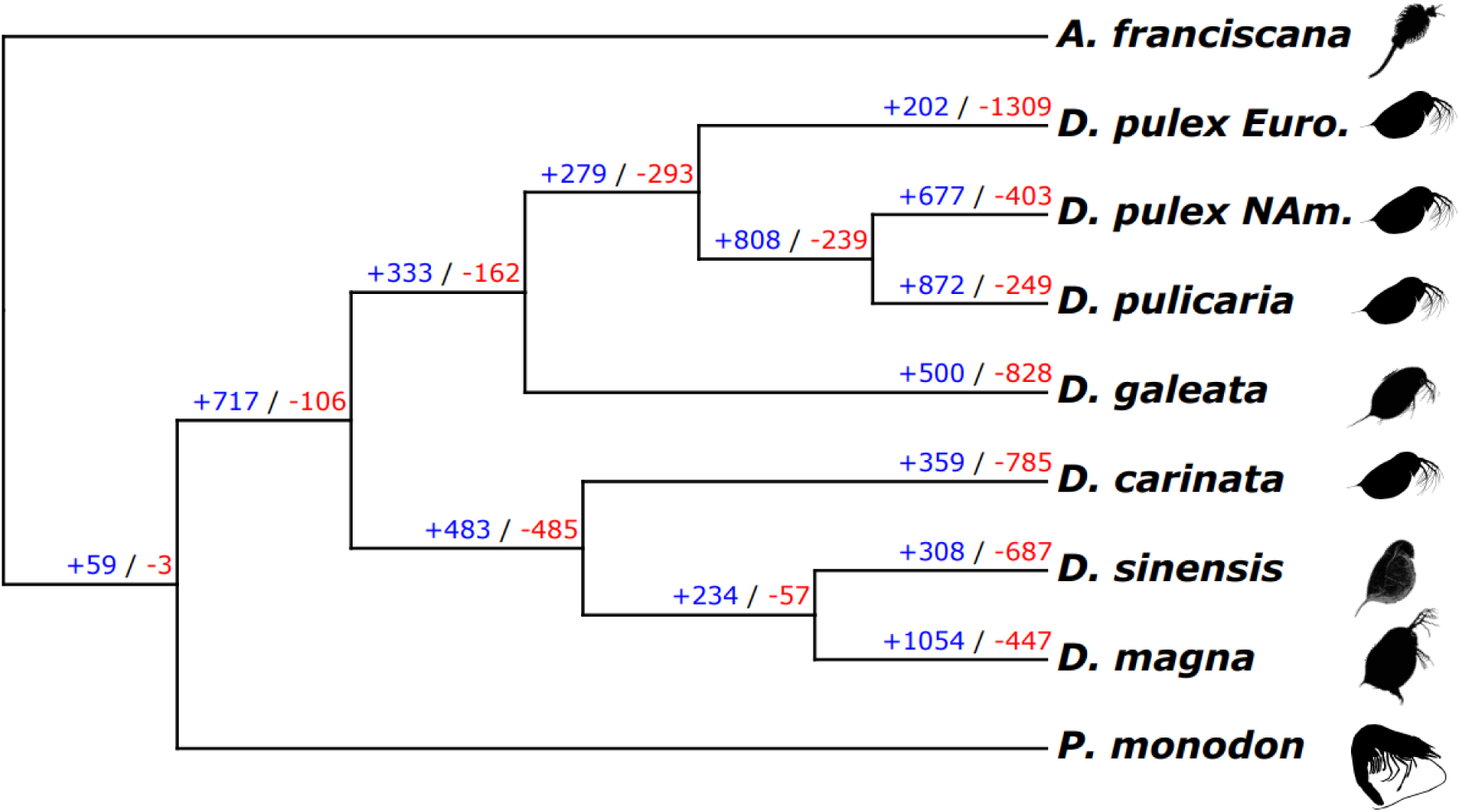
Gene family dynamics across *Daphnia.* Results of gene family evolution analyses across the phylogenetic tree from *CAFE5*. The blue colored numbers indicate the number of genes gained and the red indicate the number of genes lost within each node and terminal leaf. Icons were taken from PhyloPic.com.

Next, we were interested in understanding how gene expansions are related to function across species (Lespinet et al., 2002; Sánchez-Gracia et al., 2009), we explicitly want to understand if there are common functions across all species that could be related to ecologically relevant phenotypes for Daphnids. We examine whether there is a generality across all species within our data by first measuring the enrichment of GO terms belonging to the expanding genes within each species’ genome. We extracted the most common expanding genes in this dataset, and one intriguing pattern is that they are largely involved with glycolysis and glycoprotein biosynthetic processes (Figure 2), potentially linked with Daphnids ability to withstand periods of anoxia, food limitation, and stress response. Additionally, most Daphnids have terms related to double-strand DNA repair enriched, expect for *D. magna* and North American *D. pulex*, which may represent a general stress response to DNA damage across the genus (Figure 2). We see some general expansions for G-protein processes and phosphorylation across species as well and have a notable expansion of several terms related to immune responses in *D. magna* which is a well-known system studied for resistance to pathogens (Ebert, 2022).

**Figure 2:**
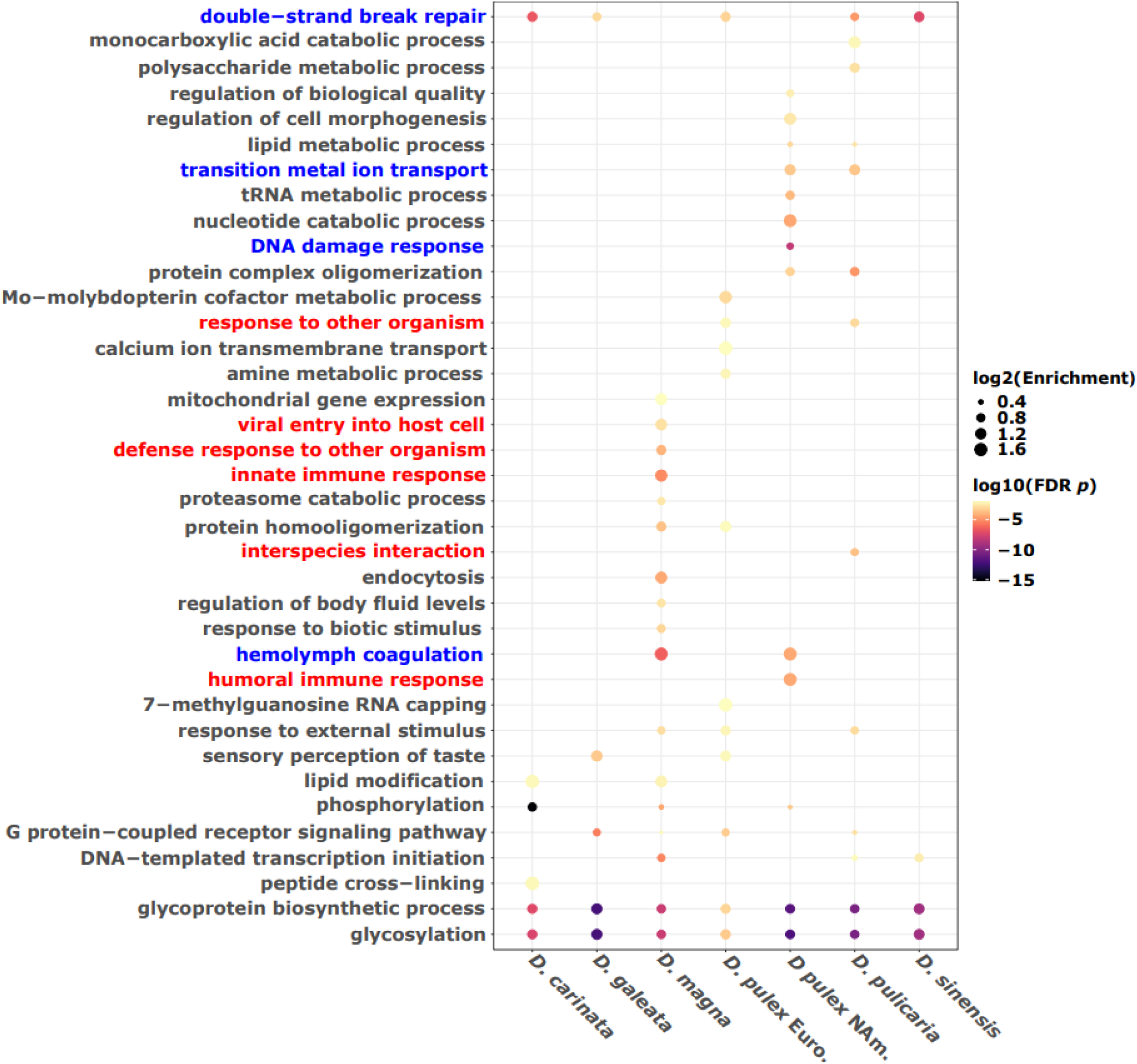
Significantly expanding gene families and their gene ontology enrichment across species reveals an excess of stress response terms. Presence and absence data of the most enriched terms across species, the y-axis is the enriched terms within each species. The red colored GO terms indicate any terms related to immune responses and the blue terms are general stress responses. All terms have been semantically reduced and condensed long descriptors.

We do not show any specific expansions of terms belonging to the *Ctenodaphnia* subgenera (e.g., *D. carinata*, *D. magna*, *D. sinensis*) save for lipid modification, however there are several terms that belong specifically to the *Daphnia* subgenera including transition metal ion transport and sensory perception of touch among others (Figure 2). Yet, there is not a specific set of terms defining the difference between subgenera (*Ctenodaphnia* vs *Daphnia*) for both contractions and expansions (Supplemental Figure 2&3).

**Supplemental Figure 2:**
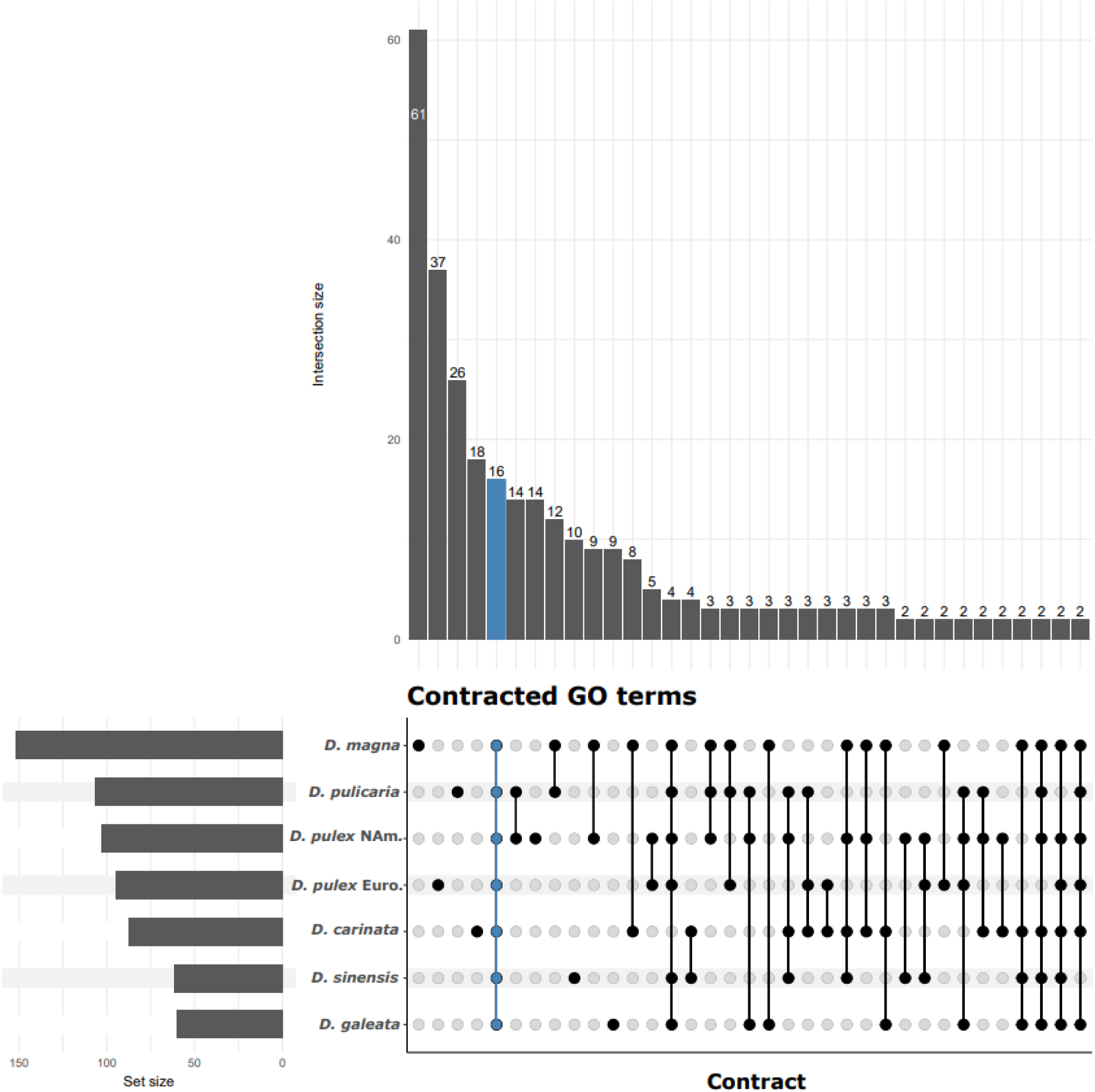
Upset plot of the enriched gene ontology terms for contracted gene families. The blue colored column denotes the GO terms shared by all species. We only show GO term combinations that appear at least twice and also remove outgroup comparisons. The insertion size (y-axis of the above barplots) is the number of GO terms shared in each group by set combination. Set size (x-axis of the side barplots) are the number of significant GO terms within each species. These terms are not semantically reduced.

**Supplemental Figure 3:**
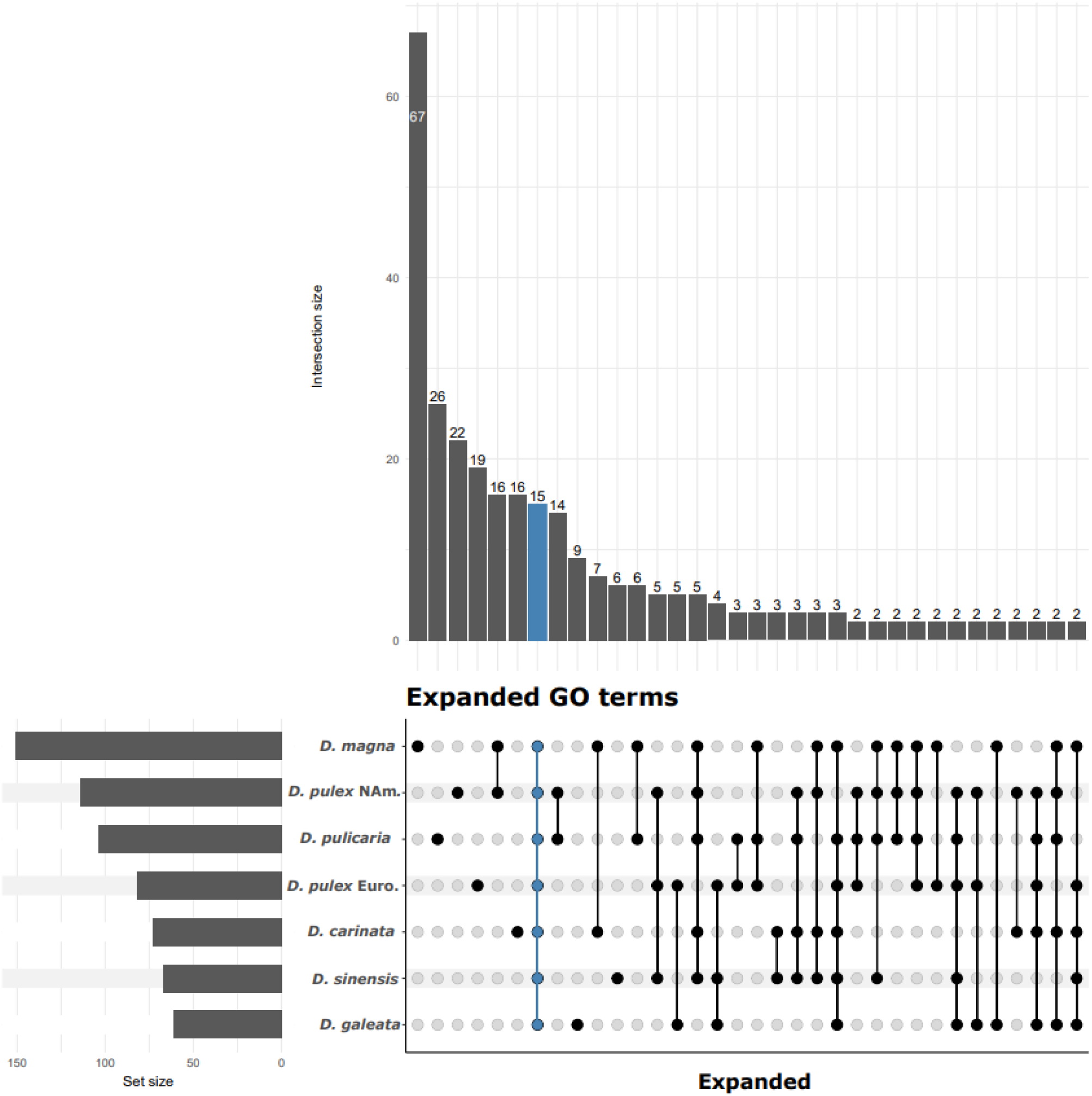
Upset plot of the enriched gene ontology terms for expanding gene families. The blue colored column denotes the GO terms shared by all species. We only show GO term combinations that appear at least twice and also remove outgroup comparisons. The insertion size (y-axis of the above barplots) is the number of GO terms shared in each group by set combination. Set size (x-axis of the side barplots) are the number of significant GO terms within each species. These terms are not semantically reduced.

Outside of these examples from the enrichment data, there are many unique expanding and contracting GO terms found to be enriched in only one or two species (Supplemental Figure 2&3). While we classify the enrichment of evolving gene families here, we test below whether these commonly enriched genes undergo selection across *Daphnia* genomes.

### General patterns of positive selection on evolving gene families

We tested the hypothesis that expanding gene families undergo higher rates of selection by using *hyphy v2.5 aBSREL* (Kosakovsky Pond et al., 2020) on codon sequences from the relevant gene families identified from *OrthoFinder*. We excluded gene families that identified extreme values of *dN/dS* (*ω*) > 10 and classified a tip, or branch, as being under positive selection if both the *dN/dS* > 1 and the multiple testing adjusted *p*-value < 0.05. We found that the expanded gene families have 6 of the 410 (1.5%) trees with positive selection compared to the non-fluctuating class (32 in 5,044; 0.63%). A Fisher’s exact test shows a positive odds ratio of 2.31 [95 % CI: 0.96, 5.6] (two-tailed Fisher’s exact test; *z* = 1.87; *p* = 0.062), yet non-significant.

### Evolution of Stress Response Gene Families and Natural Selection in Daphnia

We investigated whether gene family expansions associated with stress response GO terms identified in Figure 2 are subject to positive selection using the adaptive Branch-Site Random Effects Likelihood (*aBSREL*) method. Specifically, we examined gene families involved in iron-ion binding, heme production, and hemolymph processes due to their relevance in hypoxia adaptation and stress responses in *Daphnia* (Zeis et al., 2009). Initially, we focused on the *hemoglobin-1* gene family, which showed significant expansions within the transition metal-ion binding and hemolymph coagulation GO terms (Figure 2) and is directly involved in stress response and heme production among *Daphnia*.

Positive selection was assessed using likelihood ratio tests (LRT) with an adjusted threshold of *p* < 0.05 after multiple testing corrections. Trees were rooted using available outgroups when present. Using *aBSREL*, we identified positive selection specifically in *D. sinensis* (*p* = 0.006; Figure 3A). Despite substantial expansions in *D. galeata* (N = 5) and *D. pulicaria* (N = 7), no selection was detected in these expanded lineages (Figure 3A). We further explored the hemolymph response by examining the *2-oxoglutarate and iron-dependent oxygenase JMJD4-like* protein family, noting expansions (N = 3) in *D. magna*, though without any detectable selection. This gene family remains single-copy across all tested species and outgroups (including crustaceans *Artemia* and *Penaeus*), except *D. magna*, suggesting potential non-selective functional diversification (Supplemental Figure 4).

**Figure 3:**
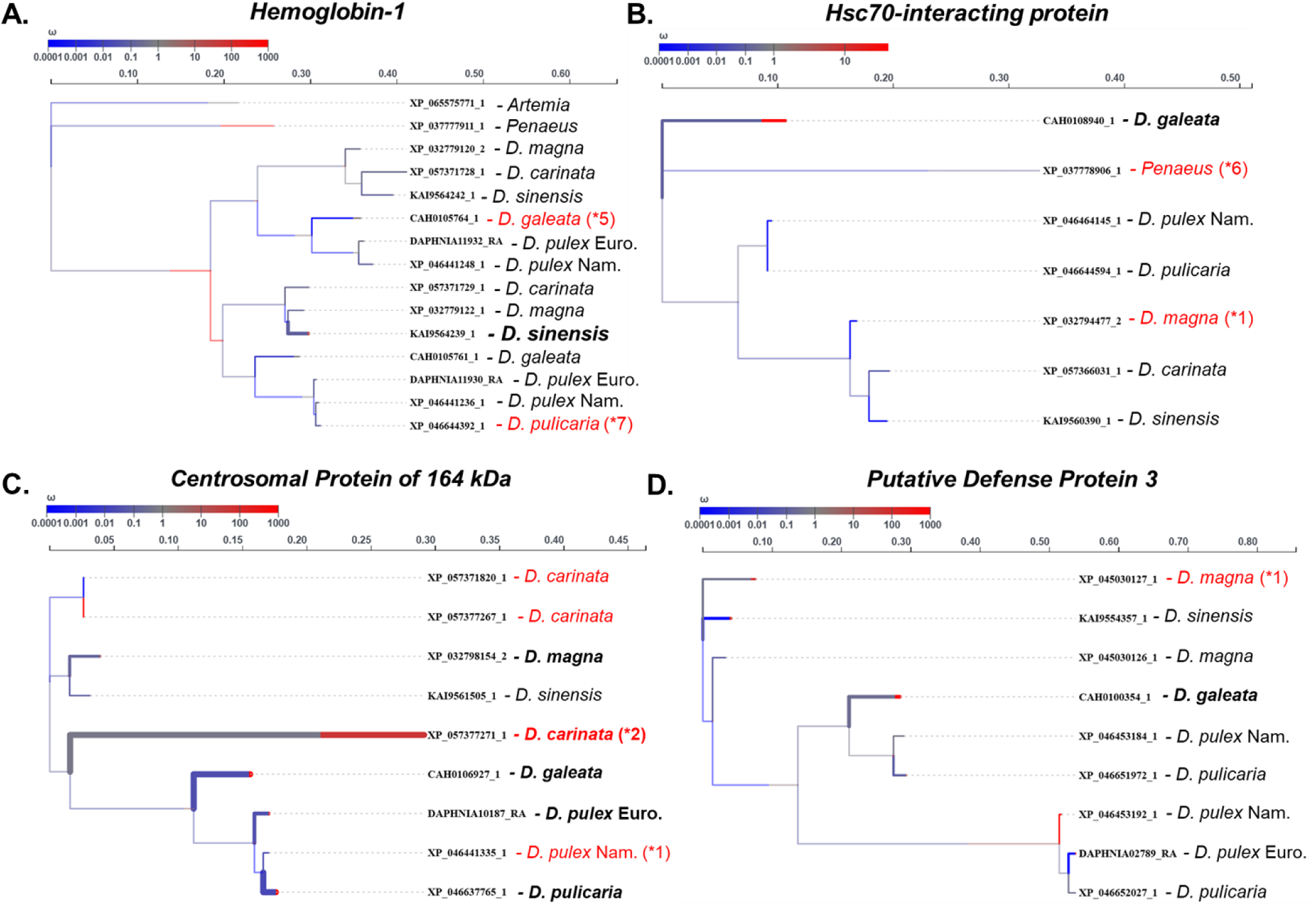
General stress response gene families undergoing positive selection and expansions. **A-D)** The thickness of branches indicates the level of significance and any tips that are significant are bolded. If a tip label is colored red, that indicates that a significant expansion occurred with the number of expanded genes in parentheses. The gene names are included as tip labels with the species name. The color of branches indicates the estimated *dN/dS*.

**Supplemental Figure 4:**
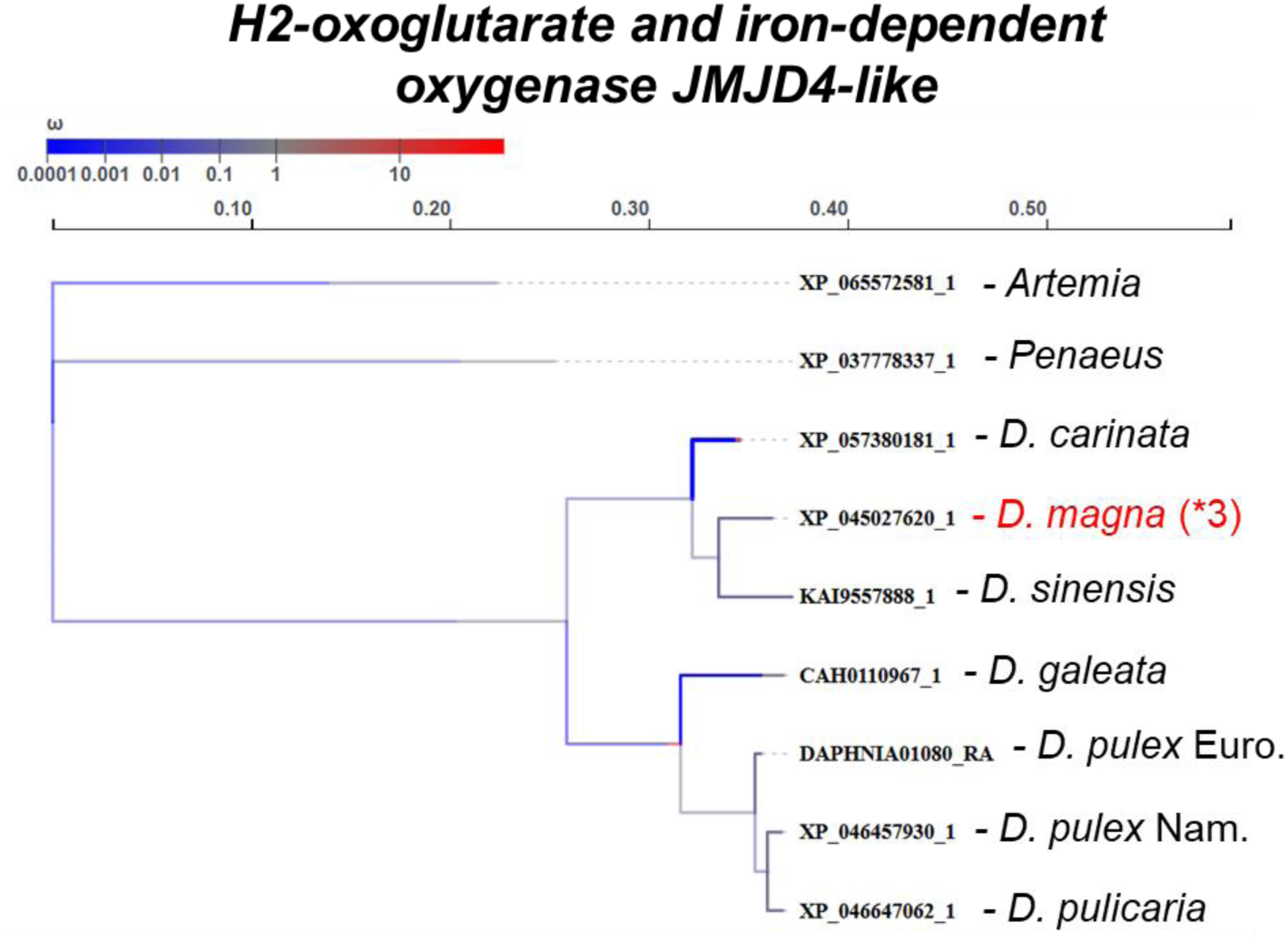
Heme response gene family with expansions, but no positive selection. The thickness of branches indicates the level of significance. If a tip label is colored red, that indicates that a significant expansion occurred with the number of expanded genes in parentheses. The gene names are included as tip labels with the species name. The color of branches indicates the estimated *dN/dS*.

Signals of selection were observed in the *hsc70-interacting* protein family within *D. galeata* (*p* = 0.046; Figure 3B), with expansions noted in both *D. magna* and the crustacean outgroup *Penaeus*. This indicates a broad evolutionary significance likely related to chaperone and heat shock protein function, which are widely conserved stress response components. Another related family, *heat shock protein beta-1-like isoform X1*, had expansions (N = 2) within *D. magna*, yet lacked detectable selection. Additionally, we explored genes related to physical defense responses in *Daphnia*, such as the *Chitin-binding type-2 domain-containing* protein family, which had expansions (N = 1) in European *D. pulex*, but showed no selection signals.

Within the DNA damage and repair pathways, the *centrosomal protein of 164 kDa-like* family exhibited selection signals across multiple branches (5 of 9 terminal branches analyzed). Specifically, *D. carinata* demonstrated strong positive selection (*p* = 1.82 × 10^−6^; Figure 3C) with gene expansions (N = 2). Interestingly, selection signals (adjusted *p* < 0.05) were widespread throughout the genus except in *D. sinensis*, and although North American *D. pulex* also exhibited expansions, no selection signals were detected. This family is likely critical for DNA repair and chromosomal organization and has broadly expanded among *Daphnia* while absent in examined outgroups (Figure 3C).

Immune-related pathways highlighted strong selection within the *clotting factor G beta subunit-like* protein family specific to North American *D. pulex* (*p* = 6.93 × 10^−12^) and associated with expansions (N = 2; Figure 2). Similarly, *putative defense protein 3*, potentially important in pathogen defense, expanded within *D. magna* (N = 1) and showed positive selection specifically in *D. galeata* (*p* = 0.0032; Figure 3D).

Reproductive gene families also showed lineage-specific dynamics, with a *testes-specific* protein family expanding and undergoing strong positive selection exclusively in *D. sinensis* (*p* = 4.92 × 10^−5^). Notably, related orthologs were restricted to *D. sinensis* and *D. magna*, suggesting specialized reproductive roles.

Analysis of glycosylation and glycoprotein synthesis terms, significantly expanded across the genus (Figure 2), revealed that the *glycoprotein-N-acetylgalactosamine 3-beta-galactosyltransferase* family underwent positive selection in *D. carinata* (*p* = 9.3 × 10^−5^) without expansions, while *D. pulicaria* exhibited both selection (*p* = 2.13 × 10^−5^) and expansion (N = 1), illustrating species-specific adaptive divergence (Supplemental Figure 5).

**Supplemental Figure 5:**
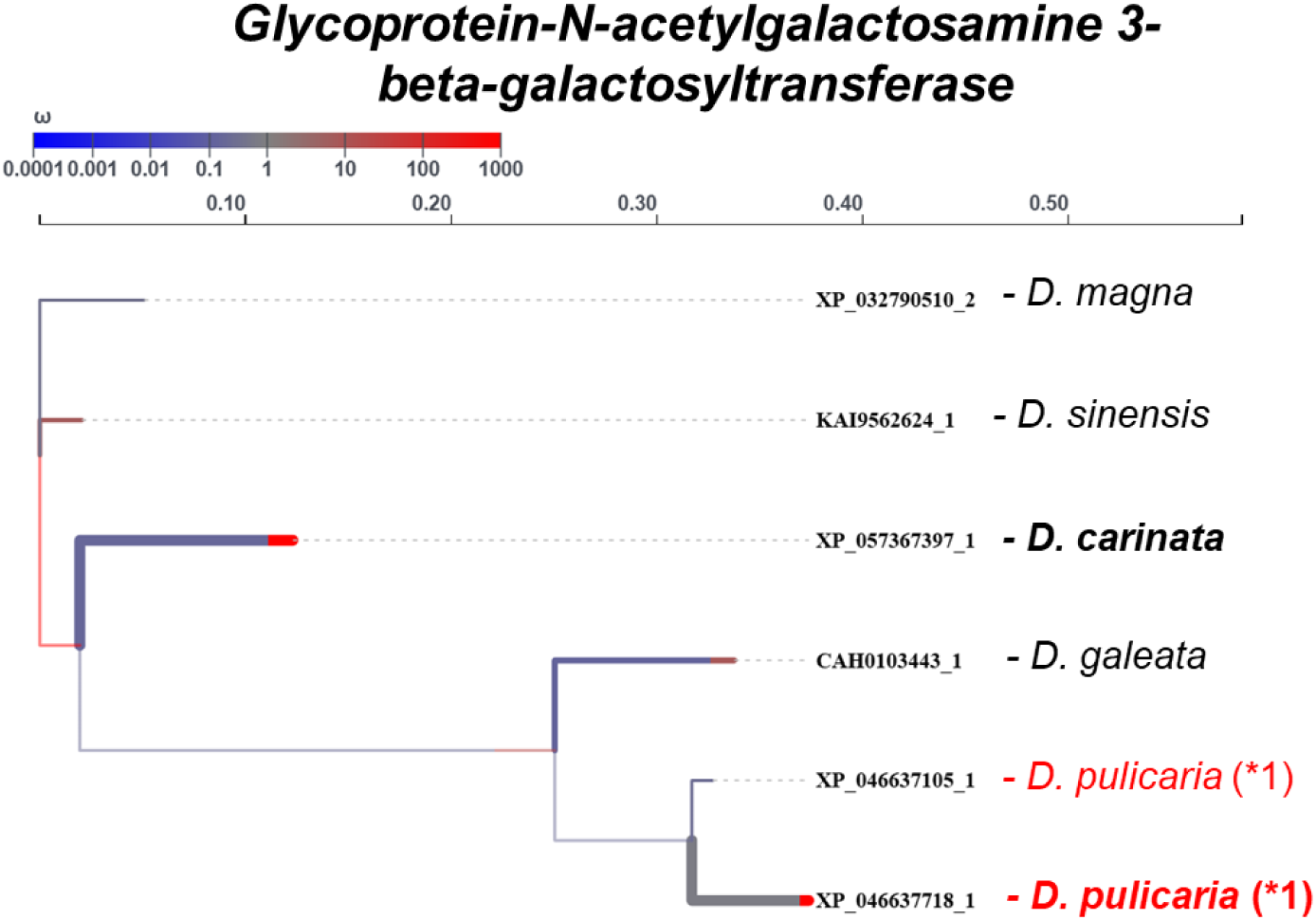
Glycoprotein synthesis gene family undergoing expansions and positive selection. The thickness of branches indicates the level of significance. If a tip label is colored red, that indicates that a significant expansion occurred with the number of expanded genes in parentheses. The gene names are included as tip labels with the species name. The color of branches indicates the estimated *dN/dS*.

Beyond specific expansions, general patterns of selection included structural and cytoskeletal GO terms related to heme production, where 6 of 133 genes (4.5%) showed selection signals. In defense and immune-related GO terms, 2 of 66 genes (3%) had detectable selection. Among reproductive genes, 1 of 60 genes (1.7%) showed selection at terminal branches. Additionally, many GO terms remained under-classified or uncharacterized through simple term matching and majority voting; among these under-classified families, 39 of 1,676 genes (2.33%) exhibited positive selection. Collectively, our findings suggest that gene family expansions combined with lineage-specific episodes of positive selection are likely crucial adaptive responses across stress response, immune defense, reproductive processes, DNA damage repair, and glycosylation pathways within the *Daphnia* genus.

## Discussion

In this work, we describe which gene families have expanded or contracted across *Daphnia*. We find overrepresented GO terms related to stress response. We show evidence for elevated positive selection across the gene families identified as being expanded across species, roughly affecting 1.5% of the expanding genes. Overall, we overview the general patterns of gene family expansion and contraction and identify candidate families under both positive selection that ultimately helps our understanding of the evolutionary dynamics occurring within *Daphnia*.

### Evolutionary dynamics of gene family evolution in Daphnia

*Daphnia* are an interesting group of taxa to study for the perspective of gene family evolution. For instance, *Daphnia* have undergone adaptive radiations, have varying modes of asexual and sexual reproduction, and phenotypic plasticity occurring within and between populations related from predator defense to male production (Chin & Cristescu, 2021; Hebert & Wilson, 1994). Yet broadly speaking, *Daphnia* are algae grazers that make up an integral part of the biomass within freshwater systems and as such, they may go through similar selection regimes related to predation (Schwartz, 1984) and or seasonal adaptation (Bergland et al., 2014; Winder et al., 2004). Our hypothesis was that *Daphnia* would have similar expansion relationships within their genomes related to these selective pressures in the wild. We show that there is a general trend towards expansions in metabolic (glycosylation and glycoprotein synthesis) and a number of stress responses in *Daphnia*, most notably DNA repair (Figure 2).

These results align in part with previous comparative analyses of *Daphnia* gene families. A comparative gene family analysis conducted by Zhang *et al*. (2021) using three *Daphnia* and several pancrustacean genomes revealed significant expansions that align with several enriched terms we identified. Particularly noteworthy were expansions related to methylation in *D. pulicaria* and North American *D. pulex*, as well as structural morphogenesis terms in *D. carinata* and North American *D. pulex*. Notably, Zhang *et al*., (2021) demonstrated that in the presence of fish kairomones, *D. mitsukuri* down-regulates terms associated with heme-production and iron binding while up-regulating those linked to chemosensory and visual perception. Therefore, it’s plausible that similar gene expression patterns could manifest in other *Daphnia* species in the presence of fish predators, a large selection pressure in ponds and lakes. Ye *et al*., (2017) conducted a gene evolution analysis of two *Daphnia* species, primarily uncovering terms related to chitin binding and oxidative stress processes. Additionally, heme production genes were shown to be highly variable in North American *D. pulex* and *D. magna*, in relation to their ability to tackle the issues of hypoxia in small ponds (Ye et al., 2017). This is interesting because *Daphnia* are known to adapt to hypoxic environments through several hemoglobin proteins (Fox et al., 1951; Kobayashi et al., 1994).

Additionally, glycosylation terms are likely composed of genes related to stress and defense response to other organisms. Previous experimental results of exposing *D. magna* to kairomones from *Triops cancriformis* (i.e., eastern tadpole shrimp) showed upexpression of chitin production and subsequent cuticle changes related to glycosylation (Otte et al., 2014). The *Daphnia* cuticle is composed of lipids and waxes, chitin, and glycosylated/unglycosylated proteins (Minelli et al., 2016). In an experimental exposure of a Chinese-derived *D. pulex* to *Microcystis*, microbes known to produce toxic algae blooms, also showed that *D. pulex* upregulate genes related to morphological change and glycoprotein synthesis (Huang et al., 2023). *Daphnia galeata* have similar responses to *Microcytis* and upregulate many overlapping terms with our expanding gene families like glycosylation and terms related to cuticle development (Kim et al., 2024). We show expansions in a glycoprotein synthesis family (Supplemental Figure 5) and evidence for selection occurring in *D. carinata* and *D. pulicaria*. It could be likely that similar proteins are actively evolving and will likely reveal mechanisms that could be indicative of specific environmental or biotic responses.

We show overlap between our results and Ye *et al*., (2017), where we primarily observe expansions in glycosylation terms, a distinction possibly stemming from our utilization of a more extensive set of ortholog groups and inclusion of additional *Daphnia* species. This underscores the value of comparative genomics in elucidating key biological processes within *Daphnia*. This brings to light that comparative genomics in *Daphnia* has led to several interesting findings. The first *Daphnia* genome, *D. pulex arenata*, a subspecies of circumarctic *D. pulex* had discovered over 30,000 unique genes (Colbourne et al., 2011). At the time, this number was over twice the amount in *Drosophila melanogaster* and humans. Upon reinvestigation, many are thought to be erroneous gene models due to fragmented draft genomes (Denton et al., 2014), however some of these erroneous genes could nonetheless be describing evolutionarily significant events or splicing variants (Ye et al., 2017). We used the error prediction feature in *CAFE5* (Han et al., 2013), which calculates a predicted influence of genome assembly error on the estimates of our gene expansion and contraction and found it to be 8%. This estimate of error is in range with other projects (Neale et al., 2017) and is similar to *Drosophila* genomes (Da Lage et al., 2019). Also, the gene family gain and loss rate (*λ*) across the phylogeny is *λ* = 0.0012, a similar estimate to projects in *Drosophila* (Da Lage et al., 2019; Hahn et al., 2007). Therefore, the genomes tested here tend to have similar evolutionary rates of gene gain and loss across Arthropods and Crustaceans. We chose to include high-quality genomes available to minimize assembly bias and we hope to include more in the future when created, especially those with chromosome-level scaffolds.

## Conclusion

Our study elucidates the gene family evolution of several members of *Daphnia,* and we provide supporting evidence that stress response genes are undergoing gene number evolution. We additionally show that some of these genes prone to expansions are also under positive selection, leading us to understand the gene diversification within *Daphnia*. Our study has important implications for continuing the work to elucidate the mechanisms that drive divergence across species, and we highlight the need to further validate how specific stress response genes are functional within species and populations of *Daphnia* (Genereux et al., 2020). Ultimately though, we began the knowledge building process necessary to link gene evolution with function across an interesting group of taxa prone to rapid adaptation.

## Author contributions

CSM: Conceptualization, Data curation, Formal analysis, Investigation, Methodology, Project administration, Software, Visualization, Writing - original draft, Writing - review & editing. AOB: Conceptualization, Funding acquisition, Project administration, Supervision, Writing - review & editing

## Acknowledgments

The authors wish to acknowledge members of the Bergland lab for their discussion and feedback related to the manuscript’s development. The authors acknowledge Research Computing at the University of Virginia for providing computational resources and technical support that have contributed to the results reported within this publication. URL: https://rc.virginia.edu.

## References

Barnard-Kubow, K. B., Becker, D., Murray, C. S., Porter, R., Gutierrez, G., Erickson, P., Nunez, J. C. B., Voss, E., Suryamohan, K., Ratan, A., Beckerman, A., & Bergland, A. O. (2022). Genetic Variation in Reproductive Investment Across an Ephemerality Gradient in *Daphnia pulex*. Molecular Biology and Evolution, 39(6), msac121. 10.1093/molbev/msac121

Bergland, A. O., Behrman, E. L., O’Brien, K. R., Schmidt, P. S., & Petrov, D. A. (2014). Genomic Evidence of Rapid and Stable Adaptive Oscillations over Seasonal Time Scales in *Drosophila*. PLoS Genetics, 10(11), e1004775. 10.1371/journal.pgen.1004775

Brandon, C. S., Greenwold, M. J., & Dudycha, J. L. (2017). Ancient and Recent Duplications Support Functional Diversity of *Daphnia* Opsins. Journal of Molecular Evolution, 84(1), 12–28. 10.1007/s00239-016-9777-1

Chang, T.-C., Yang, Y., Yasue, H., Bharti, A. K., Retzel, E. F., & Liu, W.-S. (2011). The Expansion of the PRAME Gene Family in Eutheria. PLoS ONE, 6(2), e16867. 10.1371/journal.pone.0016867

Chen, B., Feder, M. E., & Kang, L. (2018). Evolution of heat-shock protein expression underlying adaptive responses to environmental stress. Molecular Ecology, 27(15), 3040–3054. 10.1111/mec.14769

Chin, T. A., & Cristescu, M. E. (2021). Speciation in *Daphnia*. Molecular Ecology, mec.15824. 10.1111/mec.15824

Colbourne, J. K., Crease, T. J., Weider, L. J., Hebert, P. D. N., Duferesne, F., & Hobaek, A. (1998). Phylogenetics and evolution of a circumarctic species complex (Cladocera: *Daphnia pulex*). Biological Journal of the Linnean Society, 65(3), 347–365. 10.1111/j.1095-8312.1998.tb01146.x

Colbourne, J. K., Pfrender, M. E., Gilbert, D., Thomas, W. K., Tucker, A., Oakley, T. H., Tokishita, S., Aerts, A., Arnold, G. J., Basu, M. K., Bauer, D. J., Caceres, C. E., Carmel, L., Casola, C., Choi, J.-H., Detter, J. C., Dong, Q., Dusheyko, S., Eads, B. D., … Boore, J. L. (2011). The Ecoresponsive Genome of *Daphnia pulex*. Science, 331(6017), 555–561. 10.1126/science.1197761

Cornetti, L., Fields, P. D., Van Damme, K., & Ebert, D. (2019). A fossil-calibrated phylogenomic analysis of *Daphnia* and the *Daphniidae*. Molecular Phylogenetics and Evolution, 137, 250–262. 10.1016/j.ympev.2019.05.018

Crease, T. J., Omilian, A. R., Costanzo, K. S., & Taylor, D. J. (2012). Transcontinental Phylogeography of the *Daphnia pulex* Species Complex. PLoS ONE, 7(10), e46620. 10.1371/journal.pone.0046620

Da Lage, J.-L., Thomas, G. W. C., Bonneau, M., & Courtier-Orgogozo, V. (2019). Evolution of salivary glue genes in *Drosophila* species. BMC Evolutionary Biology, 19(1), 36. 10.1186/s12862-019-1364-9

Daniel, F., Revolution Analytics, & Weston, S. (2022). doMC: Foreach Parallel Adaptor for “parallel.”

Denton, J. F., Lugo-Martinez, J., Tucker, A. E., Schrider, D. R., Warren, W. C., & Hahn, M. W. (2014). Extensive Error in the Number of Genes Inferred from Draft Genome Assemblies. PLoS Computational Biology, 10(12), e1003998. 10.1371/journal.pcbi.1003998

Di Tommaso, P., Chatzou, M., Floden, E. W., Barja, P. P., Palumbo, E., & Notredame, C. (2017). Nextflow enables reproducible computational workflows. Nature Biotechnology, 35(4), 316–319. 10.1038/nbt.3820

Dos Reis, M., & Yang, Z. (2019). Bayesian Molecular Clock Dating Using Genome-Scale Datasets. In M. Anisimova (Ed.), Evolutionary Genomics (Vol. 1910, pp. 309–330). Springer New York. 10.1007/978-1-4939-9074-0_10

Dowle, M., & Srinivasan, A. (2023). data.table: Extension of “data.frame.” https://R-Datatable.Com, https://Rdatatable.Gitlab.Io/Data.Table, https://Github.Com/Rdatatable/Data.Table.

Duneau, D., Möst, M., & Ebert, D. (2022). Evolution of sperm morphology in a crustacean genus with fertilization inside an open brood pouch. Peer Community Journal, 2, e63. 10.24072/pcjournal.182

Ebert, D. (2022). *Daphnia* as a versatile model system in ecology and evolution. EvoDevo, 13(1), 16. 10.1186/s13227-022-00199-0

Emms, D. M., & Kelly, S. (2019). OrthoFinder: Phylogenetic orthology inference for comparative genomics. Genome Biology, 20(1), 238. 10.1186/s13059-019-1832-y

Forró, L., Korovchinsky, N. M., Kotov, A. A., & Petrusek, A. (2008). Global diversity of cladocerans (Cladocera; Crustacea) in freshwater. Hydrobiologia, 595(1), 177–184. 10.1007/s10750-007-9013-5

Fox, H. M., Gilchrist, B. M., and Phear, E. A. (1951). Functions of haemoglobin in *Daphnia*. Proceedings of the Royal Society of London. Series B - Biological Sciences, 138(893), 514–528. 10.1098/rspb.1951.0038

Frisch, D., Morton, P. K., Chowdhury, P. R., Culver, B. W., Colbourne, J. K., Weider, L. J., & Jeyasingh, P. D. (2014). A millennial-scale chronicle of evolutionary responses to cultural eutrophication in *Daphnia*. Ecology Letters, 17(3), 360–368. 10.1111/ele.12237

Genereux, D. P., Serres, A., Armstrong, J., Johnson, J., Marinescu, V. D., Murén, E., Juan, D., Bejerano, G., Casewell, N. R., Chemnick, L. G., Damas, J., Di Palma, F., Diekhans, M., Fiddes, I. T., Garber, M., Gladyshev, V. N., Goodman, L., Haerty, W., Houck, M. L., … Karlsson, E. K. (2020). A comparative genomics multitool for scientific discovery and conservation. Nature, 587(7833), 240–245. 10.1038/s41586-020-2876-6

Geoffrey Fryer. (1991). Functional morphology and the adaptive radiation of the *Daphniidae* (Branchiopoda: Anomopoda). Philosophical Transactions of the Royal Society of London. Series B: Biological Sciences, 331(1259), 1–99. 10.1098/rstb.1991.0001

Guijarro-Clarke, C., Holland, P. W. H., & Paps, J. (2020). Widespread patterns of gene loss in the evolution of the animal kingdom. Nature Ecology & Evolution, 4(4), 519–523. 10.1038/s41559-020-1129-2

Hahn, M. W., Han, M. V., & Han, S.-G. (2007). Gene Family Evolution across 12 *Drosophila* Genomes. PLoS Genetics, 3(11), e197. 10.1371/journal.pgen.0030197

Hamza, W., Hazzouri, K. M., Sudalaimuthuasari, N., Amiri, K. M. A., Neretina, A. N., Al Neyadi, S. E. S., & Kotov, A. A. (2023). Genome Assembly of a Relict Arabian Species of *Daphnia* O. F. Müller (Crustacea: Cladocera) Adapted to the Desert Life. International Journal of Molecular Sciences, 24(1), 889. 10.3390/ijms24010889

Han, M. V., Thomas, G. W. C., Lugo-Martinez, J., & Hahn, M. W. (2013). Estimating Gene Gain and Loss Rates in the Presence of Error in Genome Assembly and Annotation Using CAFE 3. Molecular Biology and Evolution, 30(8), 1987–1997. 10.1093/molbev/mst100

Hebert, P. D. N., & Wilson, C. C. (1994). Provincialism In Plankton: Endemism And Allopatric Speciation In Australian *Daphnia*. Evolution, 48(4), 1333–1349. 10.1111/j.1558-5646.1994.tb05317.x

Hou, Y., Sierra, R., Bassen, D., Banavali, N. K., Habura, A., Pawlowski, J., & Bowser, S. S. (2013). Molecular Evidence for β-tubulin Neofunctionalization in Retaria (Foraminifera and Radiolarians). Molecular Biology and Evolution, 30(11), 2487–2493. 10.1093/molbev/mst150

Huang, Z., Jiang, C., Gu, J., Uvizl, M., Power, S., Douglas, D., & Kacprzyk, J. (2023). Duplications of Human Longevity-Associated Genes Across Placental Mammals. Genome Biology and Evolution, 15(10), evad186. 10.1093/gbe/evad186

Huang, Y., Lu, N., Yang, T., Yang, J., Gu, L., & Yang, Z. (2023). Transcriptome analysis reveals the molecular basis of anti-predation defence in *Daphnia pulex* simultaneously responding to *Microcystis aeruginosa*. Freshwater Biology, 68(8), 1372–1385. 10.1111/fwb.14110

Jia, J., Dong, C., Han, M., Ma, S., Chen, W., Dou, J., Feng, C., & Liu, X. (2022). Multi-omics perspective on studying reproductive biology in *Daphnia sinensis*. Genomics, 114(2), 110309. 10.1016/j.ygeno.2022.110309

Jordan, I. K., Makarova, K. S., Spouge, J. L., Wolf, Y. I., & Koonin, E. V. (2001). Lineage-Specific Gene Expansions in Bacterial and Archaeal Genomes. Genome Research, 11(4), 555–565. 10.1101/gr.166001

Kalyaanamoorthy, S., Minh, B. Q., Wong, T. K. F., von Haeseler, A., & Jermiin, L. S. (2017). ModelFinder: Fast model selection for accurate phylogenetic estimates. Nature Methods, 14(6), 587–589. 10.1038/nmeth.4285

Katoh, K., & Standley, D. M. (2013). MAFFT Multiple Sequence Alignment Software Version 7: Improvements in Performance and Usability. Molecular Biology and Evolution, 30(4), 772–780. 10.1093/molbev/mst010

Kitts, P. A., Church, D. M., Thibaud-Nissen, F., Choi, J., Hem, V., Sapojnikov, V., Smith, R. G., Tatusova, T., Xiang, C., Zherikov, A., DiCuccio, M., Murphy, T. D., Pruitt, K. D., & Kimchi, A. (2016). Assembly: A resource for assembled genomes at NCBI. Nucleic Acids Research, 44(D1), D73–D80. 10.1093/nar/gkv1226

Kim, E. J., Jeon, D., Park, Y. J., Woo, H., & Eyun, S. I. (2024). Dietary exposure of the water flea *Daphnia galeata* to microcystin-LR. Animal Cells and Systems, 28(1), 25–36. 10.1080/19768354.2024.2302529

Kobayashi, M., Ishigaki, K., Kobayashi, M., Igarashi, Y., & Imai, K. (1994). Oxygen transport efficiency of multiple-component hemoglobin in *Daphnia magna*. Canadian Journal of Zoology, 72(12), 2169–2171. 10.1139/z94-289

Kosakovsky Pond, S. L., Poon, A. F. Y., Velazquez, R., Weaver, S., Hepler, N. L., Murrell, B., Shank, S. D., Magalis, B. R., Bouvier, D., Nekrutenko, A., Wisotsky, S., Spielman, S. J., Frost, S. D. W., & Muse, S. V. (2020). HyPhy 2.5—A Customizable Platform for Evolutionary Hypothesis Testing Using Phylogenies. Molecular Biology and Evolution, 37(1), 295–299. 10.1093/molbev/msz197

Kumar, S., Suleski, M., Craig, J. M., Kasprowicz, A. E., Sanderford, M., Li, M., Stecher, G., & Hedges, S. B. (2022). TimeTree 5: An Expanded Resource for Species Divergence Times. Molecular Biology and Evolution, 39(8), msac174. 10.1093/molbev/msac174

Lespinet, O., Wolf, Y. I., Koonin, E. V., & Aravind, L. (2002). The Role of Lineage-Specific Gene Family Expansion in the Evolution of Eukaryotes. Genome Research, 12(7), 1048–1059. 10.1101/gr.174302

Lewontin, R. C. (1974). The genetic basis of evolutionary change *(Vol.* 560). New York: Columbia University Press.

Lindquist, S., & Craig, E. A. (1988). The Heat-Shock Proteins. Annual Review of Genetics, 22(1), 631–677. 10.1146/annurev.ge.22.120188.003215

Lugli, G. A., Milani, C., Turroni, F., Duranti, S., Mancabelli, L., Mangifesta, M., Ferrario, C., Modesto, M., Mattarelli, P., Jiří, K., Van Sinderen, D., & Ventura, M. (2017). Comparative genomic and phylogenomic analyses of the Bifidobacteriaceae family. BMC Genomics, 18(1), 568. 10.1186/s12864-017-3955-4

Lumer, H. (1937). Growth and Maturation in the Parthenogenetic Eggs of *Daphnia magna* Strauss. CYTOLOGIA, 8(1), 1–14. 10.1508/cytologia.8.1

Lüpold, S., De Boer, R. A., Evans, J. P., Tomkins, J. L., & Fitzpatrick, J. L. (2020). How sperm competition shapes the evolution of testes and sperm: A meta-analysis. Philosophical Transactions of the Royal Society B: Biological Sciences, 375(1813), 20200064. 10.1098/rstb.2020.0064

Lynch, M. (2002). Gene Duplication and Evolution. Science, 297(5583), 945–947. 10.1126/science.1075472

Lynch, M., & Force, A. (2000). The Probability of Duplicate Gene Preservation by Subfunctionalization. Genetics, 154(1), 459–473. 10.1093/genetics/154.1.459

Lynch, M., Gutenkunst, R., Ackerman, M., Spitze, K., Ye, Z., Maruki, T., & Jia, Z. (2017). Population Genomics of *Daphnia pulex*. Genetics, 206(1), 315–332. 10.1534/genetics.116.190611

Manni, M., Berkeley, M. R., Seppey, M., Simão, F. A., & Zdobnov, E. M. (2021). BUSCO Update: Novel and Streamlined Workflows along with Broader and Deeper Phylogenetic Coverage for Scoring of Eukaryotic, Prokaryotic, and Viral Genomes. Molecular Biology and Evolution, 38(10), 4647–4654. 10.1093/molbev/msab199

Mathers, T. C., Hammond, R. L., Jenner, R. A., Hänfling, B., & Gómez, A. (2013). Multiple global radiations in tadpole shrimps challenge the concept of ‘living fossils.’ PeerJ, 1, e62. 10.7717/peerj.62

Mayr, E. (1963). Animal Species and Evolution: Harvard University Press. 10.4159/harvard.9780674865327

Mendes, F. K., Vanderpool, D., Fulton, B., & Hahn, M. W. (2021). CAFE 5 models variation in evolutionary rates among gene families. Bioinformatics, 36(22–23), 5516–5518. 10.1093/bioinformatics/btaa1022

Minelli, A., Boxshall, G., & Fusco, G. (2016). Arthropod biology and evolution. Springer-Verlag Berlin An.

Mulhair, P. O., Crowley, L., Boyes, D. H., Lewis, O. T., & Holland, P. W. H. (2023). Opsin Gene Duplication in Lepidoptera: Retrotransposition, Sex Linkage, and Gene Expression. Molecular Biology and Evolution, 40(11), msad241. 10.1093/molbev/msad241

Murray, C. S., Karram, M., Bass, D. J., Doceti, M., Becker, D., Nunez, J. C., Ratan, A., & Bergland, A. O. (2024). Trans-Specific Polymorphisms Between Cryptic *Daphnia* Species Affect Fitness and Behavior. Molecular Ecology, 34(3), e17632. 10.1111/mec.17632

Neale, D. B., McGuire, P. E., Wheeler, N. C., Stevens, K. A., Crepeau, M. W., Cardeno, C., Zimin, A. V., Puiu, D., Pertea, G. M., Sezen, U. U., Casola, C., Koralewski, T. E., Paul, R., Gonzalez-Ibeas, D., Zaman, S., Cronn, R., Yandell, M., Holt, C., Langley, C. H., … Wegrzyn, J. L. (2017). The Douglas-Fir Genome Sequence Reveals Specialization of the Photosynthetic Apparatus in Pinaceae. G3 Genes|Genomes|Genetics, 7(9), 3157–3167. 10.1534/g3.117.300078

Nickel, J., Schell, T., Holtzem, T., Thielsch, A., Dennis, S. R., Schlick-Steiner, B. C., Steiner, F. M., Möst, M., Pfenninger, M., Schwenk, K., & Cordellier, M. (2021). Hybridization Dynamics and Extensive Introgression in the *Daphnia longispina* Species Complex: New Insights from a High-Quality *Daphnia galeata* Reference Genome. Genome Biology and Evolution, 13(12), evab267. 10.1093/gbe/evab267

Novales Flamarique, I. (2013). Opsin switch reveals function of the ultraviolet cone in fish foraging. Proceedings of the Royal Society B: Biological Sciences, 280(1752), 20122490. 10.1098/rspb.2012.2490

Ohno, Susumu. (2013). Evolution by gene duplication. Springer Science & Buisness Media.

Otte, K. A., Fröhlich, T., Arnold, G. J., & Laforsch, C. (2014). Proteomic analysis of *Daphnia magna* hints at molecular pathways involved in defensive plastic responses. BMC genomics, 15, 1–17. 10.1186/1471-2164-15-306

Paul, R. J., Colmorgen, M., Pirow, R., Chen, Y.-H., & Tsai, M.-C. (1998). Systemic and metabolic responses in *Daphnia magna* to anoxia. Comparative Biochemistry and Physiology Part A: Molecular & Integrative Physiology, 120(3), 519–530. 10.1016/S1095-6433(98)10062-4

Peñalva-Arana, D. C., Lynch, M., & Robertson, H. M. (2009). The chemoreceptor genes of the waterflea *Daphnia pulex*: Many Grs but no Ors. BMC Evolutionary Biology, 9(1), 79. 10.1186/1471-2148-9-79

Puttick, M. N. (2019). MCMCtreeR: Functions to prepare MCMCtree analyses and visualize posterior ages on trees. Bioinformatics, 35(24), 5321–5322. 10.1093/bioinformatics/btz554

R Core Development Team. (n.d.). R: A language and environment for statistical computing.

Richter, D. J., Fozouni, P., Eisen, M. B., & King, N. (2018). Gene family innovation, conservation and loss on the animal stem lineage. eLife, 7, e34226. 10.7554/eLife.34226

Rivera, A. M., & Swanson, W. J. (2022). The Importance of Gene Duplication and Domain Repeat Expansion for the Function and Evolution of Fertilization Proteins. Frontiers in Cell and Developmental Biology, 10, 827454. 10.3389/fcell.2022.827454

Rouger, R., Reichel, K., Malrieu, F., Masson, J. P., & Stoeckel, S. (2016). Effects of complex life cycles on genetic diversity: Cyclical parthenogenesis. Heredity, 117(5), 336–347. 10.1038/hdy.2016.52

Saad, R., Cohanim, A. B., Kosloff, M., & Privman, E. (2018). Neofunctionalization in Ligand Binding Sites of Ant Olfactory Receptors. Genome Biology and Evolution, 10(9), 2490–2500. 10.1093/gbe/evy131

Sánchez-Gracia, A., Vieira, F. G., & Rozas, J. (2009). Molecular evolution of the major chemosensory gene families in insects. Heredity, 103(3), 208–216. 10.1038/hdy.2009.55

Sayers, E. W., Bolton, E. E., Brister, J. R., Canese, K., Chan, J., Comeau, D. C., Connor, R., Funk, K., Kelly, C., Kim, S., Madej, T., Marchler-Bauer, A., Lanczycki, C., Lathrop, S., Lu, Z., Thibaud-Nissen, F., Murphy, T., Phan, L., Skripchenko, Y., … Sherry, S. T. (2022). Database resources of the national center for biotechnology information. Nucleic Acids Research, 50(D1), D20–D26. 10.1093/nar/gkab1112

Schurko, A. M., Logsdon, J. M., & Eads, B. D. (2009). Meiosis genes in *Daphnia* pulexand the role of parthenogenesis in genome evolution. BMC Evolutionary Biology, 9(1), 78. 10.1186/1471-2148-9-78

Schwartz, S. S. (1984). Life History Strategies in *Daphnia*: A Review and Predictions. Oikos, 42(1), 114. 10.2307/3544616

Shen, W., Le, S., Li, Y., & Hu, F. (2016). SeqKit: A Cross-Platform and Ultrafast Toolkit for FASTA/Q File Manipulation. PLOS ONE, 11(10), e0163962. 10.1371/journal.pone.0163962

Smith, M. D., Wertheim, J. O., Weaver, S., Murrell, B., Scheffler, K., & Kosakovsky Pond, S. L. (2015). Less Is More: An Adaptive Branch-Site Random Effects Model for Efficient Detection of Episodic Diversifying Selection. Molecular Biology and Evolution, 32(5), 1342–1353. 10.1093/molbev/msv022

Smith, V. H., & Schindler, D. W. (2009). Eutrophication science: Where do we go from here? Trends in Ecology & Evolution, 24(4), 201–207. 10.1016/j.tree.2008.11.009

Spielman, S. J., Weaver, S., Shank, S. D., Magalis, B. R., Li, M., & Kosakovsky Pond, S. L. (2019). Evolution of Viral Genomes: Interplay Between Selection, Recombination, and Other Forces. In M. Anisimova (Ed.), Evolutionary Genomics (Vol. 1910, pp. 427–468). Springer New York. 10.1007/978-1-4939-9074-0_14

Steenwyk, J. L., Buida, T. J., Li, Y., Shen, X.-X., & Rokas, A. (2020). ClipKIT: A multiple sequence alignment trimming software for accurate phylogenomic inference. PLOS Biology, 18(12), e3001007. 10.1371/journal.pbio.3001007

Supek, F., Bošnjak, M., Škunca, N., & Šmuc, T. (2011). REVIGO Summarizes and Visualizes Long Lists of Gene Ontology Terms. PLoS ONE, 6(7), e21800. 10.1371/journal.pone.0021800

Suyama, M., Torrents, D., & Bork, P. (2006). PAL2NAL: Robust conversion of protein sequence alignments into the corresponding codon alignments. Nucleic Acids Research, 34(Web Server), W609–W612. 10.1093/nar/gkl315

Teekas, L., Sharma, S., & Vijay, N. (2022). Lineage-specific protein repeat expansions and contractions reveal malleable regions of immune genes. Genes & Immunity, 23(7), 218–234. 10.1038/s41435-022-00186-4

Thomas Lin Pedersen. (2022). patchwork: The Composer of Plots. https://Patchwork.Data-Imaginist.Com, https://Github.Com/Thomasp85/Patchwork.

Vergilino, R., Markova, S., Ventura, M., Manca, M., & Dufresne, F. (2011). Reticulate evolution of the Daphnia pulex complex as revealed by nuclear markers: Reticulate Evolution In The Daphnia Pulex Complex. Molecular Ecology, 20(6), 1191–1207. 10.1111/j.1365-294X.2011.05004.x

Villanueva, R. A. M., & Chen, Z. J. (2019). ggplot2: Elegant Graphics for Data Analysis (2nd ed.). Measurement: Interdisciplinary Research and Perspectives, 17(3), 160–167. 10.1080/15366367.2019.1565254

Wang, S., Zhang, Y., Yang, W., Shen, Y., Lin, Z., Zhang, S., & Song, G. (2023). Duplicate genes as sources for rapid adaptive evolution of sperm under environmental pollution in tree sparrow. Molecular Ecology, 32(7), 1673–1684. 10.1111/mec.16833

Wang, W., Zhang, X.-S., Wang, Z.-N., & Zhang, D.-X. (2023). Evolution and phylogenetic diversity of the aquaporin gene family in arachnids. International Journal of Biological Macromolecules, 240, 124480. 10.1016/j.ijbiomac.2023.124480

Weaver, S., Shank, S. D., Spielman, S. J., Li, M., Muse, S. V., & Kosakovsky Pond, S. L. (2018). Datamonkey 2.0: A Modern Web Application for Characterizing Selective and Other Evolutionary Processes. Molecular Biology and Evolution, 35(3), 773–777. 10.1093/molbev/msx335

Wersebe, M. J., Sherman, R. E., Jeyasingh, P. D., & Weider, L. J. (2023). The roles of recombination and selection in shaping genomic divergence in an incipient ecological species complex. Molecular Ecology, 32(6), 1478–1496. 10.1111/mec.16383

Wickham, H., Averick, M., Bryan, J., Chang, W., McGowan, L., François, R., Grolemund, G., Hayes, A., Henry, L., Hester, J., Kuhn, M., Pedersen, T., Miller, E., Bache, S., Müller, K., Ooms, J., Robinson, D., Seidel, D., Spinu, V., … Yutani, H. (2019). Welcome to the Tidyverse. Journal of Open Source Software, 4(43), 1686. 10.21105/joss.01686

Winder, M., Spaak, P., & Mooij, W. M. (2004). Trade-offs in *Daphnia* habitat selection. Ecology, 85(7), 2027–2036. 10.1890/03-3108

Wu, T., Hu, E., Xu, S., Chen, M., Guo, P., Dai, Z., Feng, T., Zhou, L., Tang, W., Zhan, L., Fu, X., Liu, S., Bo, X., & Yu, G. (2021). clusterProfiler 4.0: A universal enrichment tool for interpreting omics data. The Innovation, 2(3), 100141. 10.1016/j.xinn.2021.100141

Xu, S., Innes, D. J., Lynch, M., & Cristescu, M. E. (2013). The role of hybridization in the origin and spread of asexuality in *Daphnia*. Molecular Ecology, 22(17), 4549–4561. 10.1111/mec.12407

Xu, S., Li, L., Luo, X., Chen, M., Tang, W., Zhan, L., Dai, Z., Lam, T. T., Guan, Y., & Yu, G. (2022). *Ggtree*: A serialized data object for visualization of a phylogenetic tree and annotation data. iMeta, 1(4). 10.1002/imt2.56

Xu, S., Spitze, K., Ackerman, M. S., Ye, Z., Bright, L., Keith, N., Jackson, C. E., Shaw, J. R., & Lynch, M. (2015). Hybridization and the Origin of Contagious Asexuality in *Daphnia pulex*. *Molecular Biology and Evolution*, msv190. 10.1093/molbev/msv190

Yang, Z. (2007). PAML 4: Phylogenetic Analysis by Maximum Likelihood. Molecular Biology and Evolution, 24(8), 1586–1591. 10.1093/molbev/msm088

Ye, Z., Xu, S., Spitze, K., Asselman, J., Jiang, X., Ackerman, M. S., Lopez, J., Harker, B., Raborn, R. T., Thomas, W. K., Ramsdell, J., Pfrender, M. E., & Lynch, M. (2017). A New Reference Genome Assembly for the Microcrustacean *Daphnia pulex*. G3&#58; Genes|Genomes|Genetics, 7(5), 1405–1416. 10.1534/g3.116.038638

Zaffagnini, F., & Sabelli, B. (1972). Karyologic observations on the maturation of the summer and winter Eggs of *Daphnia pulex* and *Daphnia middendorffiana*. Chromosoma, 36(2). 10.1007/BF00285213

Zeis, B. (2020). Hemoglobin in Arthropods—*Daphnia* as a Model. In U. Hoeger & J. R. Harris (Eds.), Vertebrate and Invertebrate Respiratory Proteins, Lipoproteins and other Body Fluid Proteins (Vol. 94, pp. 163–194). Springer International Publishing. 10.1007/978-3-030-41769-7_6

Zeis, B., Lamkemeyer, T., Paul, R. J., Nunes, F., Schwerin, S., Koch, M., Schütz, W., Madlung, J., Fladerer, C., & Pirow, R. (2009). Acclimatory responses of the *Daphnia pulex* proteome to environmental changes. I. Chronic exposure to hypoxia affects the oxygen transport system and carbohydrate metabolism. BMC Physiology, 9(1), 7. 10.1186/1472-6793-9-7

Zhang, G., Fang, X., Guo, X., Li, L., Luo, R., Xu, F., Yang, P., Zhang, L., Wang, X., Qi, H., Xiong, Z., Que, H., Xie, Y., Holland, P. W. H., Paps, J., Zhu, Y., Wu, F., Chen, Y., Wang, J., … Wang, J. (2012). The oyster genome reveals stress adaptation and complexity of shell formation. Nature, 490(7418), 49–54. 10.1038/nature11413

Zhang, G., Li, C., Li, Q., Li, B., Larkin, D. M., Lee, C., Storz, J. F., Antunes, A., Greenwold, M. J., Meredith, R. W., Ödeen, A., Cui, J., Zhou, Q., Xu, L., Pan, H., Wang, Z., Jin, L., Zhang, P., Hu, H., … Froman, D. P. (2014). Comparative genomics reveals insights into avian genome evolution and adaptation. Science, 346(6215), 1311–1320. 10.1126/science.1251385

Zhang, X., Blair, D., Wolinska, J., Ma, X., Yang, W., Hu, W., & Yin, M. (2021). Genomic regions associated with adaptation to predation in *Daphnia* often include members of expanded gene families. Proceedings of the Royal Society B: Biological Sciences, 288(1955), 20210803. 10.1098/rspb.2021.0803

